# Fluctuating temperatures exacerbate the effects of nutritional stress during development in *Drosophila melanogaster*

**DOI:** 10.1101/2023.11.28.569099

**Authors:** Brooke Zanco, Juliano Morimoto, Fiona Cockerell, Christen Mirth, Carla M. Sgrò

**Affiliations:** School of Biological Sciences, Monash University, Melbourne, VIC, Australia; Division of Biosciences, University College London, London, UK; Institute of Mathematics, School of Natural and Computer Sciences, University of Aberdeen, Farser Noble Building, Aberdeen, AB24 3UE, Scotland; Programa de Pós-graduação em Ecologia e Conservação, Universidade Federal do Paraná, Curitiba, 82590-300, Brazil

**Keywords:** Temperature, Nutrition, Local Adaptation, Diel Cycles, Climate Change, Development

## Abstract

Recent studies suggest that elevated temperatures interact with nutrition to exacerbate the negative effects of poor nutrition. Furthermore, populations adapted to distinct environments differ in their sensitivity to combined thermal and nutritional stress. However, most studies have tested these effects using constant temperatures, even though animals in the wild experience daily and seasonal thermal fluctuations. Here, we used two locally-adapted populations of *D. melanogaster* from the east coast of Australia (a tropical and a temperate population) to ask whether temperature fluctuations interact with nutrition in the same manner as constant temperature conditions to shape life history traits, and how this differs across populations. We found that fluctuating temperatures exacerbate the negative effects of a poor diet when compared to constant temperatures. Moreover, the negative effects of nutritional stress were significantly greater in the tropical population. In contrast, we found that the temperate population was able to utilize nutrition in unique ways to maintain optimal viability under warmer temperatures. Our findings reveal the ways in which temperature and nutrition interact to impact key life history traits in geographically distinct populations, while also highlighting the importance of using temperature assays representative of natural diel cycles when examining insect responses to climate change.

## Introduction

Biodiversity and ecosystem dynamics are being impacted by climate change in unprecedented ways (Thomas *et al*., 2004; Dawson *et al*., 2011; Foden *et al*., 2013). Rising CO_2_ levels have not only led to an increase in mean temperatures, but accentuated variation in seasonal temperatures and precipitation. This is becoming an increasingly important source of chronic stress for animals, both directly and through complex interactions with other environmental stressors, such as the nutritional composition and availability of food (Thomas *et al*., 2004; Chevin, Lande and Mace, 2010; Foden *et al*., 2013; Long *et al*., 2014; Rosenblatt and Schmitz, 2016). These impacts are likely to vary geographically and among locally-adapted populations (Chakraborty, Sgrò and Mirth, 2020). Accurately evaluating the impacts of rapid climate change on species persistence thus requires a more holistic approach, evaluating multiple interacting environmental stressors and their effects on animal fitness simultaneously across populations.

Temperature is an important selective agent experienced by almost all life forms (Cossins and Bowler, 1987), particularly ectotherms that are highly vulnerable to shifts in the ambient thermal environment (Angilletta Jr., 2009; Clarke, 2017). While studies of populations and species sampled from along environmental gradients characterised by changes in climatic conditions provide insight into the capacity of organisms to adapt to thermal stress (Angilletta Jr., 2009; Kellermann *et al*., 2009; Liefting, Hoffmann and Ellers, 2009; Sørensen, Kristensen and Overgaard, 2016; Lasne *et al*., 2018), most studies have been performed under constant thermal regimes, even though temperature is known to vary across time and space with often pronounced diel cycles (Fischer *et al*., 2011; Colinet *et al*., 2015).

Importantly, key life history traits, such as body size and development time vary in their responses to fluctuating compared to constant thermal conditions (Brakefield and Mazzotta, 1995; Brakefield and Kesbeke, 1997; Colinet *et al*., 2007; Fischer *et al*., 2011; Carrington *et al*., 2013). For example, studies in *D. melanogaster* show that warmer temperature fluctuations reduce traits associated with fitness and cooler temperature fluctuations improve traits associated with fitness, when compared with constant temperatures of the same mean (Ketola, Kellermann, *et al*., 2012; Czarnoleski *et al*., 2013; Colinet *et al*., 2016a; Javal, Renault and Colinet, 2016). In contrast, in the mosquito *Anopheles funestus*, development rate and survival were negatively influenced by fluctuating temperature when compared to constant temperatures (Lyons, Coetzee and Chown, 2013). Thus, whether it is possible to make accurate assessments of climate vulnerability based on studies using constant thermal treatments has been questioned by many (e.g. Chown *et al*., 2009; Sgro *et al*., 2010; Fischer *et al*., 2011; Estay, Lima and Bozinovic, 2014; Ketola and Saarinen, 2015). Importantly, such assessments also ignore the fact that climate change will expose organisms to combinations of stressors.

In particular, increasing CO_2_ levels and changes in temperature and water availability under climate change are expected to alter the macronutrient content of plants and fruits in complex ways (Dietterich *et al*., 2015; Rosenblatt and Schmitz, 2016). In addition, the maturation process of fruits, from ripening to rotting, is temperature sensitive and can lead to changes in the density and diversity of yeast species upon which insects like *Drosophila* feed (Morais *et al*., 1995), resulting in additional changes in the ratio of protein to carbohydrate (P:C) in and around rotting fruit (Tournas and Katsoudas, 2005).

How ectotherms respond to climate change will depend on their response to combinations of stressors simultaneously (reviewed in Kaunisto, Ferguson and Sinclair, 2016), which may or may not reflect how they respond to single stressors in isolation (Sisodia and Singh, 2010). Temperature and nutrition are strongly linked by an organism’s metabolism and energy requirements (Rho and Lee, 2017). For instance, the capacity of ectotherms to absorb macronutrients from food sources can be highly dependent on temperature (Clissold, Coggan and Simpson, 2013; Clissold and Simpson, 2015). Thus, even when an insect feeds on a plant with the same nutritional composition, the effects of that plant on insect fitness will differ with thermal conditions (Stamp and Yang, 1996; Clissold and Simpson, 2015). As a result, changes in the nutritional and thermal environment can be expected to have unexpected impacts on key fitness-related traits in ectotherms. Recent work examining the interactive effects of temperature and nutrition have shown that temperature modulates the effects of nutrition on life history traits in often unpredictable ways (Kutz, Sgrò and Mirth, 2019; Chakraborty, Sgrò and Mirth, 2020).

Further, populations sampled from along environmental (latitudinal) gradients often vary in their responses to thermal stress due to local adaptation (Angilletta Jr., 2009; Liefting, Hoffmann and Ellers, 2009; Lasne *et al*., 2018). Because such populations are also exposed to different nutritional environments (Pärtel, Laanisto and Zobel, 2007), they might also be expected to differ in their sensitivity to nutritional stress. Indeed, Chakraborty et al. (2020) found that locally-adapted populations of *D. melanogaster* collected along the Australian east coast vary in their response to combined thermal and nutritional stress, but not in a way that was explained by their latitude of origin. This result might be explained by the studies’ use of constant temperature regimes.

Populations living at opposing ends of a latitudinal cline are subject to contrasting daily mean temperatures, but perhaps more importantly, different amplitudes of temperature fluctuations over time. As a result, temperate populations might be better adapted to thermal fluctuations than their tropical counterparts, as they experience more variable environments (Buckling *et al*., 2007; Hoffmann and Ma, 2015; Shah, Funk and Ghalambor, 2017; Nati *et al*., 2021). Given that insect responses to constant versus fluctuating thermal conditions can vary for a range of life history traits (Brakefield and Mazzotta, 1995; Brakefield and Kesbeke, 1997; Colinet *et al*., 2007; Fischer *et al*., 2011), the interactive effects of fluctuating thermal conditions and nutritional stress on geographically-distinct populations might be expected to produce unique phenotypes. Because temperate populations are adapted to more variable environments, we might predict that they will be better able to buffer the response of life history traits against combinations of stressors when temperatures fluctuate (Bradshaw, 1965; De Jong, 1995; Sørensen, Kristensen and Overgaard, 2016; Shah, Funk and Ghalambor, 2017; Nati *et al*., 2021).

Here, we first test whether locally-adapted populations differ in the response of their key life history traits to either fluctuating or constant temperature treatments in *D. melanogaster*. Specifically, we measured the effects of constant versus fluctuating temperatures during larval development on egg-to-adult viability and final body size. We then asked whether the same populations vary in their response to combined nutritional and thermal stress using fluctuating thermal conditions. We used Nutritional Geometry to generate 25 diets which varied in their protein and carbohydrate concentrations and their caloric content, as described in Chakraborty et al. (2020), across three thermal regimes. The response of each trait was then mapped onto a three-dimensional nutrient space to identify the thermal and nutritional optima for each population. Although laboratory experiments cannot capture the complexity of animal responses to climate change in nature, our experimental design allows us to better understand ectotherm responses to combined stressors. Moreover, our findings shed light on how adaptation to more variable thermal conditions may impact animal responses to fluctuating temperature regimes and how this will interact with nutrition to affect population persistence (Angilletta Jr., 2009; Chakraborty, Sgrò and Mirth, 2020; Litton *et al*., 2020).

## Methodology

### Fly stocks

Mass bred populations were created using flies collected from a tropical (Townsville, latitude:19.29S) and a temperate (Melbourne, latitude: 37.73) region along the east coast of Australia in April 2016 (Clemson, Sgrò and Telonis-Scott, 2016). Populations were maintained in numbers of approximately 1,500 flies and under a constant temperature of 25°C, with 12-hr light: 12-hr dark photoperiods on a yeast-dextrose-potato medium (potato flakes 20g/L; dextrose 30g/L; Brewer’s yeast 40 g/L; agar 7g/L; nipagen 6mL/L; and propionic acid 2.5 mL/L). These populations were maintained in the laboratory for 60 generations prior to experimental work described below.

### Nutritional Geometry

Twenty-five experimental diets varying in their protein, carbohydrate, and calorie concentrations were created. These diets were produced following the protocol from Kutz, Sgrò and Mirth (2019) and Chakraborty, Sgrò and Mirth (2020). Five different P:C ratios (1:8, 1:4, 1:3, 2:3, and 3:2) were created by varying the quantities of inactive yeast, dextrose, and potato flakes. This range of P:C ratios reflects those which are observed in rotting fruits in *D. melanogaster’s* natural environment (Matavelli *et al*., 2015; Silva-Soares *et al*., 2017). Each ratio was prepared at a concentration of roughly 620g of dry nutrient mass per litre. This was then sequentially diluted by 50% using a 0.5% agar solution, producing six different concentrations per ratio and giving a total of 25 diets.

### Experimental Temperature Regimes

Three different fluctuating temperature regimes with different average daily means were used: 18 °C (13–23 °C), 25 °C (20–30 °C) and 28 °C (23– 33 °C). These thermal regimes were chosen because they represent the temperature range experienced in the field during the warmer (summer and spring) months in temperate and tropical Australia (Van Heerwaarden, Kellermann and Sgrò, 2016; bom.gov.au).

### Experimental set-up

Before initiating each experimental block, flies from each population were left to lay eggs over a 24 h period in six 250 ml bottles containing approximately 62.5 ml of standard food medium. Flies eclosing from these bottles were used as the parental flies in all experiments. Parent flies were placed in laying pots containing the standard food medium described above with added blue food colorant and double agar (∼16%) and left to oviposit overnight at 25°C. Eggs were then collected and transferred into 7mL of treatment food, at a density of 20 eggs per vial, for each of the experimental treatments described below.

To determine the effects of fluctuating versus constant temperatures on egg-to-adult viability and wing size, for each population, 20 eggs were transferred into 20 replicate vials, all containing the standard yeast-dextrose-potato food medium (P:C 1:3 at a concentration of 320 Kcal/L). These vials were placed in a 25°C constant temperature cabinet with a 12-hr L: 12-hr D photoperiod, and egg-adult viability and body size of emerging flies compared to those flies that had developed on the same standard diet but under the 25 ± 5°C fluctuating treatment (see below). This process was completed in four experimental blocks over the course of eight weeks.

To compare trait responses to nutrition across the fluctuating thermal regimes, for each population 10 replicate vials of each of the 25 diets described above were placed into temperature cabinets set at 18, 25 or 28 ± 5°C (described above). Larvae were subsequently left to feed ad libitum and complete development, and egg-adult viability and body size of emerging flies was assessed (see below). All cabinets were maintained at a 12-hr L: 12-hr D photoperiod. Vials were repositioned within cabinets daily to avoid temperature and light gradient biases. This process was completed in six experimental blocks over the course of eight weeks.

### Egg-to-adult viability

To assess egg-to-adult viability, eclosed adults were transferred out of the experimental vials and counted. Vials were checked daily until all adults had eclosed. The assay ended when four consecutive vial checks generated no flies. At this point vials were discarded. Viability was measured by allocating each individual a score of 1 if they survived to eclosion or 0 if they died.

### Body size

Wing centroid size was used as a proxy for body size (Clemson et al 2018). The right wing was collected from 15 females for each temperature-diet treatment. Because size is a sexually dimorphic trait (Clemson, Sgrò and Telonis-Scott, 2018), only female wings were used. Once wings had been removed, they were placed on a microscope slide and photographed (Leica M80 stereo microscope; Leica, Heerbrugg, Switzerland). Wing centroid size was then measured using the software tpsDIG and CoordGen8 following Clemson et al. (2016).

### Statistical analysis

#### Constant versus fluctuating temperatures

To determine whether both traits vary in their response to constant versus fluctuating temperature independent of diet, we first restricted our analysis to data from the standard food, 25°C constant/25°C fluctuating experimental treatments. The models included thermal treatment (constant or fluctuating) and population as fixed effects, and experimental block as a random effect for viability and replicate for wing size. For all analyses of egg-to-adult viability and female wing centroid size (body size), linear mixed effects models were fitted to the data using a Binomial distribution for viability and a Gaussian distribution for female wing centroid size. To do this, the lme4 package was used along with emmeans and lsmeans, for pairwise comparisons. Box plots were made in Graphpad Prism (version 8.4.2).

#### Interactive effects of nutrition and fluctuating thermal regimes

To examine the extent to which fluctuating thermal conditions interact with nutrition to affect the traits examined, and whether this varied across locally adapted populations, we analysed the full data set of three fluctuating thermal conditions and 25 diets. The full model for both traits included the linear and quadratic components of protein (P, P^2^), carbohydrate (C, C^2^), their interaction (P x C) (Lee and Jang 2014), population (Pop), temperature (T) as fixed effects and all interaction terms for; temperature-by-nutrient (T x P, T x C, T x P x C), temperature-by-population (T x Pop), population-by-nutrient (Pop x P, Pop x C, Pop x P x C) and temperature-by-nutrient-by-population (T x P x Pop, T x C x Pop, T x, C x P x Pop) also as fixed effects. Experimental block was included as a random effect for viability and replicate for wing size (Kutz, Sgrò and Mirth, 2019; Chakraborty, Sgrò and Mirth, 2020).

Trait response surfaces were visualised using non-parametric thin-plate splines (TPS) from the fields package in R (version 3.4.1). All statistical analyses were conducted using R (Version 3.3.2) (R Core Team 2016). Microsoft Excel (version 12.3.6) (Microsoft, Redmond) was used for data entry and management.

#### Nutritional trade-offs

We used the newly developed ‘Nutrigonometry’ model to test for nutritional trade-offs emerging as a result of fluctuating thermal regimes across the trait response surfaces (Morimoto *et al*., 2023). Briefly, Nutrigonometry uses triangulation to identify the peak (or valley) regions of the trait response surfaces, from which comparisons can be made by applying Pythagoras theorem (Morimoto *et al*., 2023). Two metrics are calculated using Nutrigonometry: (a) the angle θ_*i,j*_, which is an estimate of the distance between the identified location of peak regions in two response surfaces for traits *i* and *j*. The metric θ_*i,j*_, therefore is a proxy for nutritional trade-offs with respect to the ratio of nutrients. (b) the hypothenuse *h_i,j_*, which is an estimate of the distance between the identified peak regions with respect to each other and to the origin. The metric *h_i,j_* therefore is a proxy for nutritional trade-offs with respect to nutrient intake or quantity (details in Morimoto *et al*., 2023). We used Nutrigonometry with a general linear model specification as it was shown to lead to the most accurate identification of peak regions in trait response surfaces (Morimoto *et al*., 2023). We estimated the peak region as the region comprising the highest 90% values for the trait response and these regions are represented as red polygons in our trait response surfaces. For each trait response (i.e., viability and wing size), we compared the estimates of θ_*i,j*_ and *h_i,j_* between populations (Melbourne versus Townsville) and amongst temperatures (18 ± 5°C, 25 ± 5°C, 28 ± 5°C) using the confidence interval of the differences between these metrics (see Supplementary Table 4).

## Results

### Fluctuating versus constant temperatures

Changes in trait responses under fluctuating versus constant temperatures have been well documented across a broad range of taxa including *Drosophila melanogaster*, *Petrolisthes cinctipes*, *Petrolisthes eriomerus, and Mytilus edulis* (Brakefield and Mazzotta, 1995; Brakefield and Kesbeke, 1997; Colinet *et al*., 2007; Bozinovic *et al*., 2011; Fischer *et al*., 2011). We also know that responses to temperature vary among geographically distinct populations (Angilletta Jr., 2009; Liefting, Hoffmann and Ellers, 2009; Kellermann, Van Heerwaarden and Sgrò, 2017). As a result, we first measured whether egg-to-adult viability and final body size differed under fluctuating versus constant temperature assays in the temperate (Melbourne) and tropical (Townsville) populations used in our study.

We compared viability and wing size measurements for flies developed at a constant temperature of 25°C, on our standard lab food, with those developed at 25 ± 5°C, fed the equivalent diet (P:C 1:3 at a concentration of 320 Kcal/L). We found that egg-to-adult viability was significantly reduced in flies reared under fluctuating temperatures when compared to a constant temperature (Figure 1a, Supplementary Table 1). The direction of this effect was consistent among populations (Figure 1a). In contrast, there was no significant effect of fluctuating temperatures on body size within individual populations (Figure 1b, Supplementary Table 2), however, when reared under fluctuating temperatures the Melbourne population was significantly larger than the Townsville population (Figure 1b). These findings indicate that fluctuating temperature treatments impact fitness-related traits both in a different manner to constant temperatures and in a population dependent manner.

**Figure 1.**
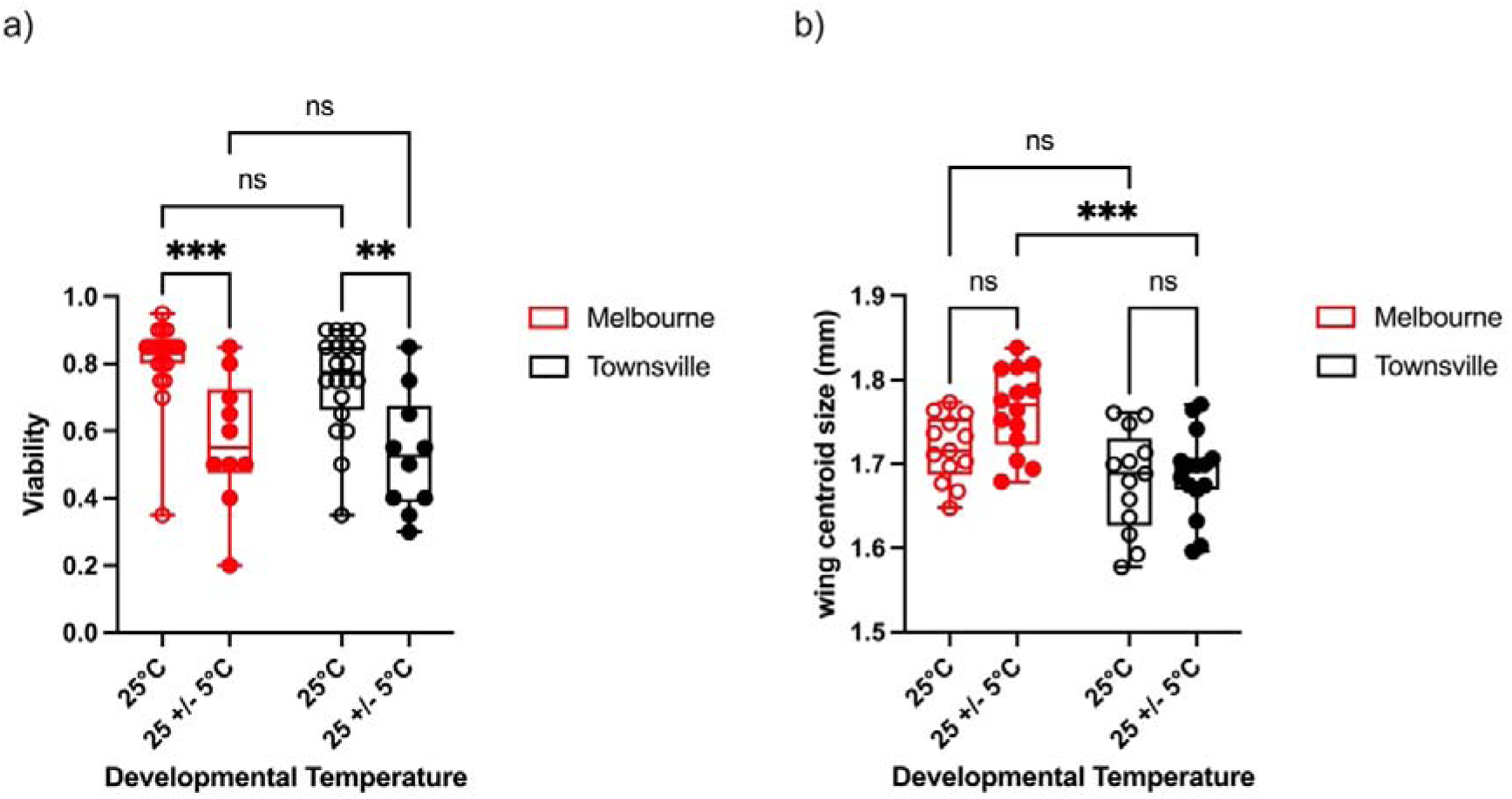
Fluctuating temperature versus constant temperature regimes modify egg-to-adult viability (a) and wing centroid size (b) in significantly different ways in populations of flies from Melbourne or Townsville. The direction of this variation differs between traits, with flies reared under fluctuating temperatures being significantly less viable when compared with those reared under constant temperatures (a). Alternatively, there was no significant effect of fluctuating temperature on body size within populations (b), however, under fluctuating temperature, body size was significantly reduced in the Townsville population when compared with the Melbourne population (b). Pairwise comparisons are shown above (black asterisks denote level of significance). All other statistical analyses are reported in Supplementary Tables 1 and 2.

### Egg-adult viability at fluctuating temperatures

Next, we aimed to uncover how temperature treatments that fluctuated around different means mediate egg-to-adult viability and body size responses to nutrition across two locally adapted populations. Egg-to-adult viability was measured across 25 experimental diets (varying in their protein: carbohydrate concentration and caloric content), three temperature treatments (18, 25 and 28 ± 5°C), and two populations (temperate (Melbourne) and tropical (Townsville)). The Melbourne population, which is adapted to more extreme thermal fluctuations, showed higher egg-to-adult viability across all three temperature treatments when compared to the Townsville population (Figure 2, Supplementary Table 3). Importantly, there was a four-way interaction between carbohydrate, protein, population, and temperature (Supplementary Table 3). As a result, we used the newly developed ‘Nutrigonometry’ model to test for complex nutritional trade-offs in both populations (Morimoto *et al*., 2023).

**Figure 2.**
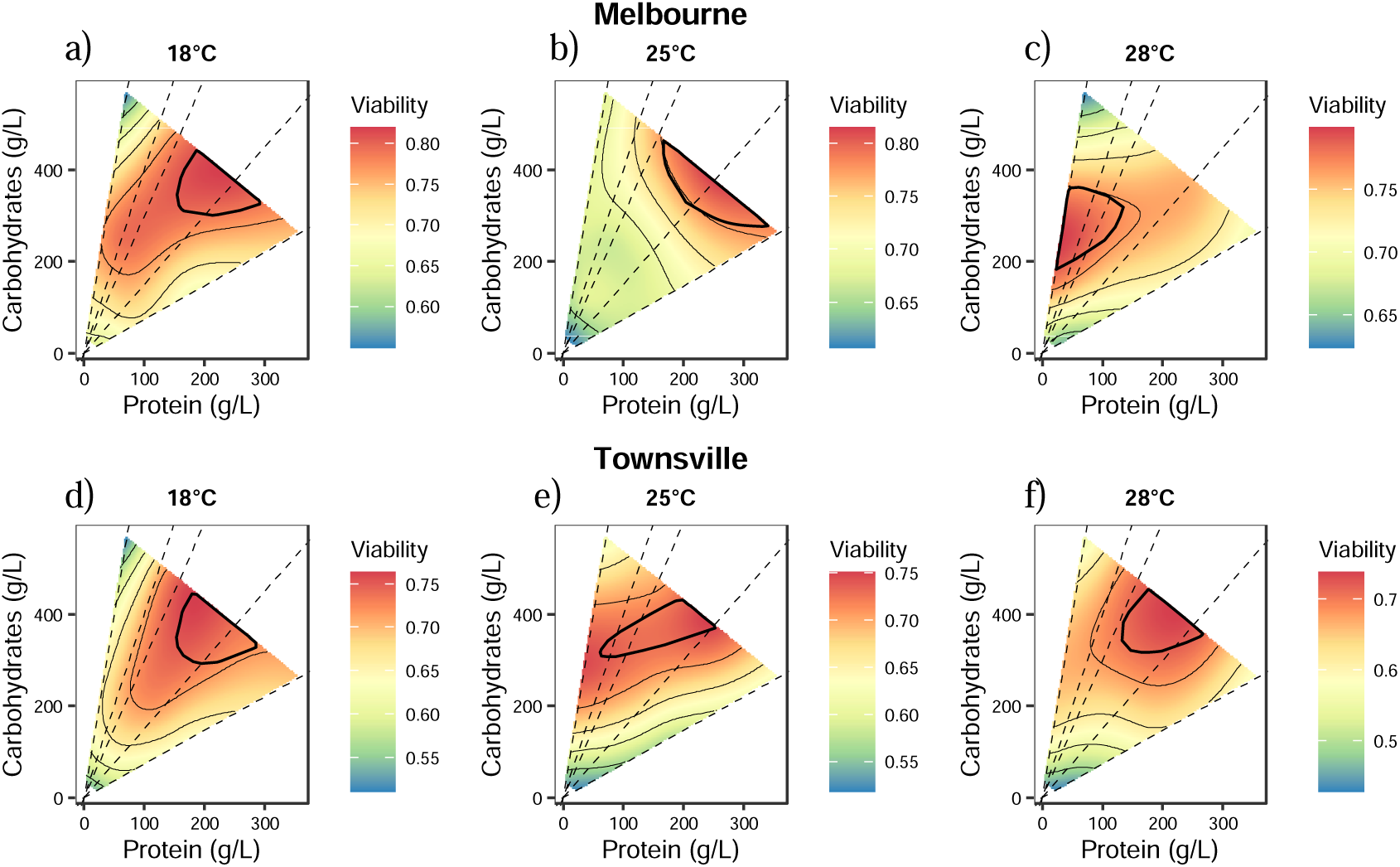
Changes in egg-to-adult viability in response to protein and carbohydrate content of the larval diet across three fluctuating temperature treatments: 18±5°C (panels a and d), 25±5°C (panels b and e), and 28±5°C (panels c and f). In all panels dashed black lines represent protein: carbohydrate ratios. Red circles represent the nutritional optimum which is calculated using the Nutrigonometry model (see Methods for details). Panels a-c show response surfaces for the Melbourne population, and panels d-e show response surfaces for the Townsville population. Statistical analysis reported in Supplementary Table 4.

The dietary conditions that generated the highest viability shifted substantially towards lower protein: carbohydrate ratios (i.e., more carbohydrate) as mean temperatures increased in the Melbourne population (Figure 2a – c, Supplementary Table 4). For instance, the peak region for viability shifted from an intermediate protein: carbohydrate ratio and high nutrient concentration at 18°C± 5°C and 25°C± 5°C to relatively lower protein: carbohydrate ratio and lower nutrient concentration at 28°C (Figure 2a-c, Supplementary Table 4). This allowed the Melbourne population to maintain relatively high viability despite thermally stressful conditions. For the Townsville population, the dietary conditions which generated the highest egg-to-adult viability remained at an intermediate-high protein: carbohydrate ratio and high nutrient concentration across thermal regimes (Figure 2 d-f, Supplementary Table 4). These findings suggest that locally-adapted populations might be expected to utilise nutrients in different ways under thermal stress. This is curious, given a previous study using constant temperature treatments did not observe this response (Chakraborty, Sgrò and Mirth, 2020).

### Body size at fluctuating temperatures

To determine the combined effects of nutrition and fluctuating temperature on final body size among two locally-adapted populations, we measured female wing size as a proxy for body size, using viable adults from the experiment described above. Overall, body size decreased with increasing rearing temperatures in both populations (see Figure 3), while the Melbourne population remained consistently larger than the Townsville population for each temperature regime (Figure 3, Supplementary Table 5). As was the case with viability, we found a four-way interaction between carbohydrate, protein, population, and temperature (Supplementary Table 5), so we again applied the ‘Nutrigonometry’ model to test for complex nutritional trade-offs in both populations (Morimoto *et al*., 2023).

**Figure 3.**
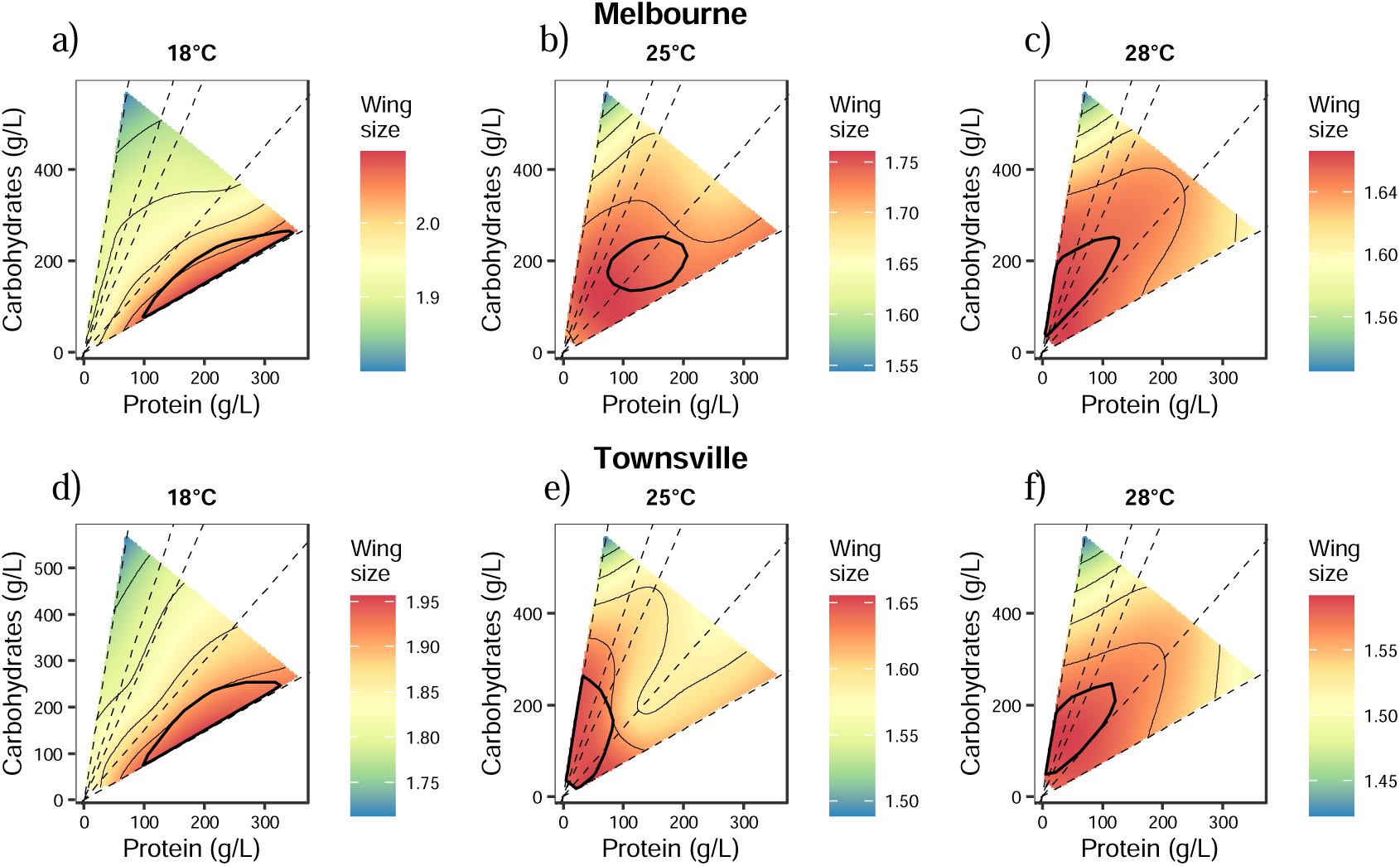
Changes in adult wing area in response to protein and carbohydrate content of the larval diet across three fluctuating temperature treatments - 18±5°C (panels a and d), 25±5°C (panels b and e), and 28±5°C (panels c and f). In all panels dashed black lines represent protein: carbohydrate ratios. Red circles represent the nutritional optimum which is calculated using the Nutritional Geometry model (see Methods for details). Panels a-c show response surfaces for the Melbourne population, and panels d-e show response surfaces for the Melbourne population. Each panel has a different scale to account for the decline in wing size as temperatures increase along with variation among populations. Statistical analysis reported in Supplementary Table 4.

There was a shift of the peak region for size as temperature increased, going from regions of high protein: carbohydrate ratios and intermediate nutrient concentration to regions of lower protein: carbohydrate ratios and low nutrient concentration in both populations (Figure 3). In the Melbourne population, there was a clear intermediate stage, where the peak in the body size surface was at an intermediate protein: carbohydrate ratio as temperature moved from 18 ±5°C to 25 ±5°C, before shifting to low protein: carbohydrate ratio at 28±5°C (Figure 3 a – c, Supplementary Table 4). This intermediate stage was not observed in the Townsville population, for which the peak in the body size surface shifted rapidly from high to low protein: carbohydrate as temperatures increased from 18±5°C to 25±5°C (Figure 3 d-f, Supplementary Table 4). The peak remained at low protein: carbohydrate ratio in the Townsville population at 28±5°C (Figure 3).

## Discussion

When considering the effects of climate change on animal fitness we must consider the effects of various environmental factors which act both independently and in combination to either alter the effects of each. Specifically, we know that both temperature and the nutritional composition and abundance of primary producers are expected to change significantly as climate change worsens. These two stressors have also been shown to interact in complex ways, with nutrition mediating the effects of temperature on key life history traits in small ectotherms (Clissold and Simpson, 2015; Kutz, Sgrò and Mirth, 2019; Chakraborty, Sgrò and Mirth, 2020; Kim, Jang and Lee, 2020).

While previous studies provide insights into how temperature changes responses to diet, they have used constant temperature assays, which may over or underestimate the impacts of increasing temperatures on animal fitness (Ketola *et al*., 2012; Czarnoleski *et al*., 2013; Colinet *et al*., 2016; Javal, Renault and Colinet, 2016). Constant temperature regimes also fail to consider how local adaptation to thermal variability in nature may impact the ability of populations to tolerate the increasing temperatures (and increases in thermal fluctuations) expected under climate change. To these ends, the first aim of this study was to understand how key life history traits (viability and final body size) respond to the effects of fluctuating versus constant temperatures during development among two locally-adapted populations of *D. melanogaster* collected from along a latitudinal cline in eastern Australia. The second aim of this study was to investigate the interactive effects of fluctuating temperatures and nutrition on the same two traits.

### Populations vary in their response to constant versus fluctuating temperature

Our findings for viability are generally in line with previous investigations which show that key life history traits are negatively impacted by fluctuating warm versus constant warm temperatures (Ketola, Kristensen, *et al*., 2012; Colinet *et al*., 2016; Javal, Renault and Colinet, 2016). We found that in both populations, viability was significantly reduced under fluctuating temperature treatments, when compared with constant temperature treatments. Our results suggest that phenotypic plasticity in egg-to-adult viability differs in response to fluctuating versus constant temperature regimes. This may be because thermal oscillations more closely mimic developmental conditions in the wild (Les, Paitz and Bowden, 2009). It is also possible that constant temperatures have different effects on an animal’s ability to regulate their metabolism than fluctuating temperatures (Lamb, 1961). In the wild, animals will typically decrease their metabolic rate in response to thermal stress – this is done by reducing their activity and feeding (Barnes and Laurie-Ahlberg, 1986; Klepsatel et al., 2019; Vianna *et al*., 2020). This process may be disrupted under constant temperature conditions. These findings highlight the need for fluctuating temperature treatments to be increasingly included in studies of population responses to environmental change.

Interestingly, we found that final body size was not significantly reduced when flies were developed at fluctuating temperatures versus constant temperatures. This is interesting in light of findings by Czarnoleski *et al. (*2013), who reported a reduction in *D. melanogaster* wing size when exposed to fluctuating temperatures during development. Similar to our study, flies were reared at two mean temperatures (18 and 25°C), either constant or ±4°C (Czarnoleski *et al*., 2013). However, it is important to note that the population used in Czarnoleski *et al*., 2013 was collected from a site in the USA, thus, it is possible that these contrasting findings represent variability in phenotypic responses among different populations.

Indeed, we found a significant difference among populations for body size, but only when flies were reared under fluctuating temperatures. In this instance, the temperate population was significantly larger than the tropical population at fluctuating compared to constant temperatures. We know that populations sampled from latitudinal gradients adapt to their local climatic conditions and that the temperate Melbourne population is subject to more severe and more frequent thermal shifts (Hoffmann et al. 2001, 2002, 2003; Lasne et al. 2018; Sgrò et al. 2010). They are also likely to be adapted to a more variable nutritional environment, given these shifts in temperature (Bradshaw, 1965; De Jong, 1995; Sørensen, Kristensen and Overgaard, 2016). This leads us to believe that populations adapted to greater variation in thermal maxima and minima, and to greater nutritional variability can maintain larger body sizes under more stressful (in this case fluctuating) thermal conditions. This supports the environmental variability hypothesis which predicts that populations which are adapted to less variable environments (tropical populations) should display reduced plasticity when compared with populations subjected to more variable environmental conditions (temperate populations) (Bradshaw, 1965; Stearns and Kawecki, 1994; De Jong, 1995; Wagner, Booth and Bagheri-Chaichian, 1997).

### Local adaptation changes population responses to combined thermal and nutritional stress

As a result of local adaptation, stress traits associated with starvation resistance in *Drosophila* often vary across latitudinal clines (Sisodia and Singh, 2010; Lasne *et al*., 2018). Thus, population differences in trait values found along latitudinal gradients might be driven by the combined effects of changing nutritional and thermal conditions (Robinson and Partridge, 2001; Kolss *et al*., 2009; Flatt, 2016). Indeed, we found that nutrition mediates the effects of fluctuating temperatures on both egg-to-adult viability and body size, and that these responses are population specific.

As discussed previously, egg-to-adult viability is often considered a canalized trait in ectotherms, and this trait has previously been found not to vary significantly among populations, steadily declining as temperatures increase (Conover and Schultz, 1995; Wagner, Booth and Bagheri-Chaichian, 1997; Kutz, Sgrò and Mirth, 2019; Chakraborty, Sgrò and Mirth, 2020). Interestingly, we found that the temperate population was able to maintain on average higher egg-to-adult viability than the tropical population and that under warmer temperatures this effect was mediated by the nutritional environment. Specifically, the temperate population appeared to dramatically shift its nutritional optimum from an intermediate P:C diet towards a low protein, high carbohydrate diet, whereas in the tropical population the same diet produced maximum viability across all thermal conditions. We know that different traits have different macronutrient requirements for maximal performance, and that this can vary in different thermal environments (Lee *et al*., 2008; Boggs, 2009). However, our findings suggest that larvae from our temperate population can utilise their nutritional environment in unique ways to maintain optimal viability under a range of thermal conditions.

Interestingly, a study by Chakraborty, Sgrò and Mirth (2020) found no interactive effect of temperature and nutrition on egg-to-adult viability among different locally adapted populations – however, they used constant 18 and 25°C temperature assays. Similarly, the caterpillar *Spodoptera exigua* (Hübner) had the highest viability at intermediate P:C ratios (Lee and Roh, 2010), regardless of whether they were reared at 18, 26, or 34 °C. These findings suggest that under constant temperatures the interactive effects of temperature and nutrition are dampened when compared to the effects under fluctuating thermal conditions.

Previous work has shown that a protein rich diet maximises body size in *Drosophila* (Kristensen *et al*., 2011; Rodrigues *et al*., 2015; Poças, Crosbie and Mirth, 2020). Our results at 18 ± 5°C align with this existing knowledge. Interestingly however, at 28 ± 5°C both populations had larger wings when reared on low protein, low carbohydrate concentration diets. This suggests that wing size does not increase with high protein at warmer temperatures, as it does at lower temperature. Furthermore, given these results, we hypothesise that viability/body size trade-offs, whereby the nutritional optima for each trait diverges, may become more extreme under warmer conditions, particularly for the tropical population, where high protein diets maximised viability but reduced body size at 28± 5°C.

Under warmer mean temperatures, metabolic stress has been shown to increase in ectotherms (Rho and Lee, 2017) and is modulated by developmental nutrition. For example, when reared at warmer temperatures (28 °C), animals on low-protein diets showed the highest metabolic rates (Alton *et al*., 2020). As a result, it is unsurprising that we observed narrower viability and body size peaks at 28 ± 5°C compared to 18 and 25± 5°C, in both populations. This means that the number of diets which generate the optimal phenotype are reduced under increasing thermal conditions. Similar findings were reported by Kutz et al. (2019) using constant temperature treatments. This suggests that this reduction in nutritional optima is not specific to fluctuating temperature regimes but observed under higher temperatures in general. This has significant implications for ectotherm persistence under climate change, as it would be expected to significantly reduce the ecological niche of different populations.

Our findings indicate that it might not be possible to determine a universal predictive model or explanation for nutrient by temperature interactions in ectotherms. We know that nutrition and temperature are linked via an organism’s metabolism and energy requirements (Cross *et al*., 2015; Clissold and Simpson, 2015). Thus, given that plant species distribution and abundance is influenced by climate (reviewed in Rosenblatt and Schmitz, 2016), it may be that locally-adapted populations have evolved different means of moderating thermal stress via diet, based on the resources available to them. This hints at the possibility that the populations examined in this study may have different physiological means of utilising macronutrients, particularly under warmer conditions. For instance, they might vary in how they process food through digestion, absorb nutrients, or store excess energy (Volkoff and Rønnestad, 2020).

### Responses to environmental change are complex – broader implications, limitations, and future directions

In this study, we established that egg-adult viability and wing size differ in their responses to constant versus fluctuating temperatures in *D. melanogaster*. Then, using Nutritional Geometry, we found that the response of these traits to the combined effects of nutrition and temperature was population specific. Even within a single population, trait responses to nutrition differed with temperature treatments, producing unexpected results under warmer conditions. This suggests that the ways in which locally-adapted populations respond to multiple simultaneous drivers of global change are likely to differ significantly.

Nevertheless, it must be noted that there were several limitations to our study. First, the way protein increase was achieved in the yeast-dextrose-potato media was to increase the yeast component. However, yeast also contains other essential macro and micronutrients, and these micronutrients might interact with protein and carbohydrates to dictate life-history outcomes (Rodrigues *et al*., 2015; Zanco *et al*., 2021; Piper *et al*., 2022). Therefore, the life-history traits examined in this study may not only be responding to carbohydrate and protein content, but also to complex interactions between these other nutritional elements. Nevertheless, we know that the range of P:C ratios used in the current study do reflect those which are observed in rotting fruits in the natural environment (Matavelli *et al*., 2015; Silva-Soares *et al*., 2017). This indicates that laboratory-based experiments such as ours still play an important role in understanding the responses of ectotherms to multiple simultaneous drivers of global change.

Another limitation of our study was the fact that temperature fluctuations typically experienced at lower latitudes are less severe than those experienced in more temperate environments (Cunningham and Read, 2003). Thus, the reduced viability of the tropical population across all treatments could be attributed to the fact that the extremity of fluctuations used in the current study were more pronounced than the temperature fluctuations tropical populations would experience in nature. Exploring how traits respond to differing amplitudes of thermal fluctuations would be an interesting avenue of future research. Adding to this, it would also be beneficial for future research to determine which fruits our study populations feed on in the wild and investigate how different thermal conditions influence macronutrient ratios during the decomposition processes. This would provide invaluable insights into what might be driving variation in trait responses to nutrition under different thermal conditions observed in this study.

Finally, given our results suggest that different populations may have different physiological mechanisms of dealing with thermal stress via nutrition, it would be useful to establish what molecular pathways may be mediating these differences. This would allow us to better understand the mechanisms that underpin differences among locally-adapted populations, and further our understanding of the ways in which different populations might evolve in response to ongoing environmental change.

## Conclusions

Environmental change is not one-dimensional, instead it is the result of multiple interacting factors, varying over space and time. Despite this, most studies aimed at better understanding and predicting species responses to environmental change focus on static temperatures and single stressors within single populations. Our study shows that locally-adapted populations respond to the combined effects of nutrition and fluctuating thermal regimes in trait and population-specific ways. Even within single populations, we found that trait responses to nutrition varied across temperature treatments.

In conclusion, our study suggests that the effects of fluctuating temperature are mediated by diet and that these two environmental factors interact to shape fitness outcomes, making it difficult to predict the way traits will respond with increasing temperatures. Furthermore, the effects of these two stressors on larval traits are population specific. Therefore, creating predictive models about the effects of climate change on organisms may only be possible at more local scales, and with nutrition and temperature considered as interacting factors.

## Author contributions

**Brooke Zanco:** Conceptualization, Data curation, Formal analysis, Investigation, Visualization, Methodology, Writing - original draft

**Juliano Morimoto:** Formal analysis, Visualization, Methodology, Writing – review and editing

**Fiona Cockerell:** Investigation

**Christen Mirth:** Conceptualization, Formal analysis, Supervision, Funding acquisition, Methodology, Project administration, Writing - review and editing

**Carla M. Sgrò:** Conceptualization, Formal analysis, Supervision, Funding acquisition, Methodology, Project administration, Writing - review and editing

## Funding and Acknowledgements

This research was supported by funds from the Australian Research Council, DP180103725 to CMS and FT170100259 to CKM, and the School of Biological Sciences, Monash University. JM is supported by the BBSRC (BB/V015249/1).

## Data availability

The data and scripts underlying this article are available at: DOI: 10.26180/24616122

## References

Alton, L. A. et al. (2020) ‘Developmental nutrition modulates metabolic responses to projected climate change’, Functional Ecology, 34(12), pp. 2488–2502. doi: 10.1111/1365-2435.13663.

Angilletta Jr., M. J. (2009) Thermal Adaptation. Oxford University Press. doi: 10.1093/acprof:oso/9780198570875.001.1.

Barnes, C. P. and Laurie-Ahlberg, C. C. (1986) ‘Genetic Variability of Flight Metabolism I N’, Genetics, pp. 267–294.

Boggs, C. L. (2009) ‘Understanding insect life histories and senescence through a resource allocation lens’, Functional Ecology, 23(1), pp. 27–37. doi: 10.1111/j.1365-2435.2009.01527.x.

Bozinovic, F. et al. (2011) ‘The mean and variance of environmental temperature interact to determine physiological tolerance and fitness’, Physiological and Biochemical Zoology, 84(6), pp. 543–552. doi: 10.1086/662551.

Bradshaw, A. D. (1965) ‘Evolutionary Significance of Phenotypic Plasticity in Plants’, in, pp. 115–155. doi: 10.1016/S0065-2660(08)60048-6.

Brakefield, P. M. and Kesbeke, F. (1997) ‘Genotype–environment interactions for insect growth in constant and fluctuating temperature regimes’, Proceedings of the Royal Society of London. Series B: Biological Sciences, 264(1382), pp. 717–723. doi: 10.1098/rspb.1997.0102.

Brakefield, P. M. and Mazzotta, V. (1995) ‘Matching field and laboratory environments: effects of neglecting daily temperature variation on insect reaction norms’, Journal of Evolutionary Biology, 8(5), pp. 559–573. doi: 10.1046/j.1420-9101.1995.8050559.x.

Buckling, A. et al. (2007) ‘Experimental adaptation to high and low quality environments under different scales of temporal variation’, Journal of Evolutionary Biology, 20(1), pp. 296–300. doi: 10.1111/j.1420-9101.2006.01195.x.

Carrington, L. B. et al. (2013) ‘Effects of Fluctuating Daily Temperatures at Critical Thermal Extremes on Aedes aegypti Life-History Traits’, PLoS ONE, 8(3). doi: 10.1371/journal.pone.0058824.

Chakraborty, A., Sgrò, C. M. and Mirth, C. K. (2020) ‘Does local adaptation along a latitudinal cline shape plastic responses to combined thermal and nutritional stress?’, Evolution, 74(9), pp. 2073–2087. doi: 10.1111/evo.14065.

Chevin, L.-M., Lande, R. and Mace, G. M. (2010) ‘Adaptation, Plasticity, and Extinction in a Changing Environment: Towards a Predictive Theory’, PLoS Biology. Edited by J. G. Kingsolver, 8(4), p. e1000357. doi: 10.1371/journal.pbio.1000357.

Chown, S. L. et al. (2009) ‘Phenotypic variance, plasticity and heritability estimates of critical thermal limits depend on methodological context’, Functional Ecology, 23(1), pp. 133–140. doi: 10.1111/j.1365-2435.2008.01481.x.

Clarke, A. (2017) Princliples of Thermal Ecology. Oxford University Press.

Clemson, A. S., Sgrò, C. M. and Telonis-Scott, M. (2016) ‘Thermal plasticity in Drosophila melanogaster populations from eastern Australia: quantitative traits to transcripts’, Journal of Evolutionary Biology, 29(12), pp. 2447–2463. doi: 10.1111/jeb.12969.

Clemson, A. S., Sgrò, C. M. and Telonis-Scott, M. (2018) ‘Transcriptional profiles of plasticity for desiccation stress in Drosophila’, Comparative Biochemistry and Physiology Part B: Biochemistry and Molecular Biology, 216, pp. 1–9. doi: 10.1016/j.cbpb.2017.11.003.

Clissold, F. J., Coggan, N. and Simpson, S. J. (2013) ‘Insect herbivores can choose microclimates to achieve nutritional homeostasis’, Journal of Experimental Biology. doi: 10.1242/jeb.078782.

Clissold, Fiona J and Simpson, S. J. (2015) ‘Temperature, food quality and life history traits of herbivorous insects’, Current Opinion in Insect Science, 11, pp. 63–70. doi: 10.1016/j.cois.2015.10.011.

Clissold, Fiona J. and Simpson, S. J. (2015) ‘Temperature, food quality and life history traits of herbivorous insects’, Current Opinion in Insect Science. Elsevier Inc, 11, pp. 63–70. doi: 10.1016/j.cois.2015.10.011.

Colinet, H. et al. (2007) ‘Proteomic profiling of a parasitic wasp exposed to constant and fluctuating cold exposure’, Insect Biochemistry and Molecular Biology, 37(11), pp. 1177–1188. doi: 10.1016/j.ibmb.2007.07.004.

Colinet, H. et al. (2015) ‘Insects in Fluctuating Thermal Environments’, Annual Review of Entomology, 60(1), pp. 123–140. doi: 10.1146/annurev-ento-010814-021017.

Colinet, H. et al. (2016a) ‘Uncovering the benefits of fluctuating thermal regimes on cold tolerance of drosophila flies by combined metabolomic and lipidomic approach’, Biochimica et Biophysica Acta (BBA) - Molecular and Cell Biology of Lipids, 1861(11), pp. 1736–1745. doi: 10.1016/j.bbalip.2016.08.008.

Colinet, H. et al. (2016b) ‘Uncovering the benefits of fluctuating thermal regimes on cold tolerance of drosophila flies by combined metabolomic and lipidomic approach’, Biochimica et Biophysica Acta - Molecular and Cell Biology of Lipids, 1861(11), pp. 1736–1745. doi: 10.1016/j.bbalip.2016.08.008.

Conover, D. O. and Schultz, E. T. (1995) ‘Phenotypic similarity and the evolutionary significance of countergradient variation’, Trends in Ecology & Evolution, 10(6), pp. 248–252. doi: 10.1016/S0169-5347(00)89081-3.

Cossins, A. R. and Bowler, K. (1987) Temperature Biology of Animals. 1st edn. Springer Dordrecht.

Cross, W. F. et al. (2015) ‘Interactions between temperature and nutrients across levels of ecological organization’, Global Change Biology, 21(3), pp. 1025–1040. doi: 10.1111/gcb.12809.

Cunningham, S. C. and Read, J. (2003) ‘Do temperate rainforest trees have a greater ability to acclimate to changing temperatures than tropical rainforest trees?’, New Phytologist, 157(1), pp. 55–64. doi: 10.1046/j.1469-8137.2003.00652.x.

Czarnoleski, M. et al. (2013) ‘Flies developed small bodies and small cells in warm and in thermally fluctuating environments’, Journal of Experimental Biology, 216(15), pp. 2896–2901. doi: 10.1242/jeb.083535.

Dawson, T. P. et al. (2011) ‘Beyond Predictions: Biodiversity Conservation in a Changing Climate’, Science, 332(6025), pp. 53–58. doi: 10.1126/science.1200303.

Dietterich, L. H. et al. (2015) ‘Impacts of elevated atmospheric CO2 on nutrient content of important food crops’, Scientific Data, 2(1), p. 150036. doi: 10.1038/sdata.2015.36.

Estay, S. A., Lima, M. and Bozinovic, F. (2014) ‘The role of temperature variability on insect performance and population dynamics in a warming world’, Oikos, 123(2), pp. 131–140. doi: 10.1111/j.1600-0706.2013.00607.x.

Fischer, K. et al. (2011) ‘Assay conditions in laboratory experiments: Is the use of constant rather than fluctuating temperatures justified when investigating temperature-induced plasticity?’, Oecologia, 166(1), pp. 23–33. doi: 10.1007/s00442-011-1917-0.

Flatt, T. (2016) ‘Genomics of clinal variation in Drosophila: disentangling the interactions of selection and demography’, Molecular Ecology, 25(5), pp. 1023–1026. doi: 10.1111/mec.13534.

Foden, W. B. et al. (2013) ‘Identifying the World’s Most Climate Change Vulnerable Species: A Systematic Trait-Based Assessment of all Birds, Amphibians and Corals’, PLoS ONE. Edited by S. Lavergne, 8(6), p. e65427. doi: 10.1371/journal.pone.0065427.

van Heerwaarden, B., Kellermann, V. and Sgrò, C. M. (2016) ‘Limited scope for plasticity to increase upper thermal limits’, Functional Ecology, 30(12), pp. 1947–1956. doi: 10.1111/1365-2435.12687.

Javal, M., Renault, D. and Colinet, H. (2016) ‘Impact of fluctuating thermal regimes on Drosophila melanogaster survival to cold stress’, Animal Biology, 66(3–4), pp. 427–444. doi: 10.1163/15707563-00002510.

De Jong, G. (1995) ‘Phenotypic Plasticity as a Product of Selection in a Variable Environment’, The American Naturalist, 145(4), pp. 493–512. Available at: https://www.jstor.org/stable/2462965.

Kaunisto, S., Ferguson, L. V and Sinclair, B. J. (2016) ‘Can we predict the effects of multiple stressors on insects in a changing climate?’, Current Opinion in Insect Science, 17, pp. 55–61. doi: 10.1016/j.cois.2016.07.001.

Kellermann, V. et al. (2009) ‘Fundamental Evolutionary Limits in Ecological Traits Drive Drosophila Species Distributions’, Science, 325(5945), pp. 1244–1246. doi: 10.1126/science.1175443.

Kellermann, V., Van Heerwaarden, B. and Sgrò, C. M. (2017) ‘How important is thermal history? Evidence for lasting effects of developmental temperature on upper thermal limits in Drosophila melanogaster’, Proceedings of the Royal Society B: Biological Sciences, 284(1855), p. 20170447. doi: 10.1098/rspb.2017.0447.

Ketola, T., Kristensen, T. N., et al. (2012) ‘Can evolution of sexual dimorphism be triggered by developmental temperatures?’, Journal of Evolutionary Biology, 25(5), pp. 847–855. doi: 10.1111/j.1420-9101.2012.02475.x.

Ketola, T., Kellermann, V., et al. (2012) ‘Constant, cycling, hot and cold thermal environments: strong effects on mean viability but not on genetic estimates’, Journal of Evolutionary Biology, 25(6), pp. 1209–1215. doi: 10.1111/j.1420-9101.2012.02513.x.

Ketola, T. and Saarinen, K. (2015) ‘Experimental evolution in fluctuating environments: tolerance measurements at constant temperatures incorrectly predict the ability to tolerate fluctuating temperatures’, Journal of Evolutionary Biology, 28(4), pp. 800–806. doi: 10.1111/jeb.12606.

Kim, K. E., Jang, T. and Lee, K. P. (2020) ‘Combined effects of temperature and macronutrient balance on life-history traits in Drosophila melanogaster: implications for life-history trade-offs and fundamental niche’, Oecologia. Springer Berlin Heidelberg, 193(2), pp. 299–309. doi: 10.1007/s00442-020-04666-0.

Klepsatel, P., Wildridge, D. and Gáliková, M. (2019) ‘Temperature induces changes in Drosophila energy stores’, Scientific Reports, 9(1), pp. 1–10. doi: 10.1038/s41598-019-41754-5.

Kolss, M. et al. (2009) ‘LIFE-HISTORY CONSEQUENCES OF ADAPTATION TO LARVAL NUTRITIONAL STRESS IN DROSOPHILA’, Evolution, 63(9), pp. 2389–2401. doi: 10.1111/j.1558-5646.2009.00718.x.

Kristensen, T. N. et al. (2011) ‘Dietary protein content affects evolution for body size, body fat and viability in Drosophila melanogaster’, Biology Letters, 7(2), pp. 269–272. doi: 10.1098/rsbl.2010.0872.

Kutz, T. C., Sgrò, C. M. and Mirth, C. K. (2019) ‘Interacting with change: Diet mediates how larvae respond to their thermal environment’, Functional Ecology, 33(10), pp. 1940–1951. doi: 10.1111/1365-2435.13414.

Lamb, K. P. (1961) ‘Some Effects of Fluctuating Temperatures on Metabolism, Development, and Rate of Population Growth in the Cabbage Aphid, Brevicoryne Brassicae Author ( s ): K. P. Lamb Published by: Wiley on behalf of the Ecological Society of America Stable URL: h’, Ecology, 42(4), pp. 740–745.

Lasne, C. et al. (2018) ‘Cross-sex genetic correlations and the evolution of sex-specific local adaptation: Insights from classical trait clines in Drosophila melanogaster’, Evolution, 72(6), pp. 1317–1327. doi: 10.1111/evo.13494.

Lee, K. P. et al. (2008) ‘Lifespan and reproduction in Drosophila: New insights from nutritional geometry’, Proceedings of the National Academy of Sciences of the United States of America, 105(7), pp. 2498–2503. doi: 10.1073/pnas.0710787105.

Lee, K. P. and Roh, C. (2010) ‘Temperature-by-nutrient interactions affecting growth rate in an insect ectotherm’, Entomologia Experimentalis et Applicata, 136(2), pp. 151–163. doi: 10.1111/j.1570-7458.2010.01018.x.

Les, H. L., Paitz, R. T. and Bowden, R. M. (2009) ‘Living at extremes: Development at the edges of viable temperature under constant and fluctuating conditions’, Physiological and Biochemical Zoology, 82(2), pp. 105–112. doi: 10.1086/590263.

Liefting, M., Hoffmann, A. A. and Ellers, J. (2009) ‘Plasticity versus environmental canalization: Population differences in thermal responses along a latitudinal gradient in drosophila serrata’, Evolution, 63(8), pp. 1954–1963. doi: 10.1111/j.1558-5646.2009.00683.x.

Litton, C. M. et al. (2020) ‘Impact of Mean Annual Temperature on Nutrient Availability in a Tropical Montane Wet Forest’, Frontiers in Plant Science, 11(June), pp. 1–13. doi: 10.3389/fpls.2020.00784.

Long, R. A. et al. (2014) ‘Behavior and nutritional condition buffer a large-bodied endotherm against direct and indirect effects of climate’, Ecological Monographs, 84(3), pp. 513–532. doi: 10.1890/13-1273.1.

Lyons, C. L., Coetzee, M. and Chown, S. L. (2013) ‘Stable and fluctuating temperature effects on the development rate and survival of two malaria vectors, Anopheles arabiensis and Anopheles funestus’, Parasites and Vectors, 6(1), pp. 1–9. doi: 10.1186/1756-3305-6-104.

Ma, G., Hoffmann, A. A. and Ma, C.-S. (2015) ‘Daily temperature extremes play an important role in predicting thermal effects’, Journal of Experimental Biology. doi: 10.1242/jeb.122127.

Ma, G., Hoffmann, A. A. and Ma, C.-S. (2021) ‘Are extreme high temperatures at low or high latitudes more likely to inhibit the population growth of a globally distributed aphid?’, Journal of Thermal Biology, 98, p. 102936. doi: 10.1016/j.jtherbio.2021.102936.

Matavelli, C. et al. (2015) ‘Differences in larval nutritional requirements and female oviposition preference reflect the order of fruit colonization of Zaprionus indianus and Drosophila simulans’, Journal of Insect Physiology, 82, pp. 66–74. doi: 10.1016/j.jinsphys.2015.09.003.

Morais, P. B. et al. (1995) ‘Yeast succession in the amazon fruit Parahancornia amapa as resource partitioning among Drosophila spp.’, Applied and Environmental Microbiology, 61(12), pp. 4251–4257. doi: 10.1128/aem.61.12.4251-4257.1995.

Morimoto, J. et al. (2023) ‘Nutrigonometry I: Using Right-Angle Triangles to Quantify Nutritional Trade-Offs in Performance Landscapes’, The American Naturalist, 201(5), pp. 725–740. doi: 10.1086/723599.

Nati, J. J. H. et al. (2021) ‘Intraspecific variation in thermal tolerance differs between tropical and temperate fishes’, Scientific Reports, 11(1), p. 21272. doi: 10.1038/s41598-021-00695-8.

Ostrowski, D. et al. (2018) ‘A biphasic locomotor response to acute unsignaled high temperature exposure in Drosophila’, PLoS ONE, 13(6), pp. 1–11. doi: 10.1371/journal.pone.0198702.

Pärtel, M., Laanisto, L. and Zobel, M. (2007) ‘CONTRASTING PLANT PRODUCTIVITY– DIVERSITY RELATIONSHIPS ACROSS LATITUDE: THE ROLE OF EVOLUTIONARY HISTORY’, Ecology, 88(5), pp. 1091–1097. doi: 10.1890/06-0997.

Piper, M. D. W. et al. (2022) ‘Dietary restriction and lifespan: adaptive reallocation or somatic sacrifice?’, FEBS Journal, pp. 1–10. doi: 10.1111/febs.16463.

Poças, G. M., Crosbie, A. E. and Mirth, C. K. (2020) ‘When does diet matter? The roles of larval and adult nutrition in regulating adult size traits in Drosophila melanogaster’, Journal of Insect Physiology. doi: 10.1016/j.jinsphys.2020.104051.

Rho, M. S. and Lee, K. P. (2017) ‘Temperature-driven plasticity in nutrient use and preference in an ectotherm’, Oecologia, 185(3), pp. 401–413. doi: 10.1007/s00442-017-3959-4.

Robinson, S. J. W. and Partridge, L. (2001) ‘Temperature and clinal variation in larval growth efficiency in Drosophila melanogaster’, Journal of Evolutionary Biology, 14(1), pp. 14–21. doi: 10.1046/j.1420-9101.2001.00259.x.

Rodrigues, M. A. et al. (2015) ‘Drosophila melanogaster larvae make nutritional choices that minimize developmental time’, Journal of Insect Physiology. Elsevier Ltd, 81, pp. 69–80. doi: 10.1016/j.jinsphys.2015.07.002.

Rosenblatt, A. E. and Schmitz, O. J. (2016) ‘Climate Change, Nutrition, and Bottom-Up and Top-Down Food Web Processes’, Trends in Ecology & Evolution, 31(12), pp. 965–975. doi: 10.1016/j.tree.2016.09.009.

Sgro, C. M. et al. (2010) ‘A comprehensive assessment of geographic variation in heat tolerance and hardening capacity in populations of Drosophila melanogaster from eastern Australia’, Journal of Evolutionary Biology, 23(11), pp. 2484–2493. doi: 10.1111/j.1420-9101.2010.02110.x.

Shah, A. A., Funk, W. C. and Ghalambor, C. K. (2017) ‘Thermal Acclimation Ability Varies in Temperate and Tropical Aquatic Insects from Different Elevations’, Integrative and Comparative Biology, 57(5), pp. 977–987. doi: 10.1093/icb/icx101.

Silva-Soares, N. F. et al. (2017) ‘Adaptation to new nutritional environments: larval performance, foraging decisions, and adult oviposition choices in Drosophila suzukii’, BMC Ecology, 17(1), p. 21. doi: 10.1186/s12898-017-0131-2.

Sisodia, S. and Singh, B. N. (2010) ‘Resistance to environmental stress in Drosophila ananassae: latitudinal variation and adaptation among populations’, Journal of Evolutionary Biology, 23(9), pp. 1979–1988. doi: 10.1111/j.1420-9101.2010.02061.x.

Sørensen, J. G., Kristensen, T. N. and Overgaard, J. (2016) ‘Evolutionary and ecological patterns of thermal acclimation capacity in Drosophila: is it important for keeping up with climate change?’, Current Opinion in Insect Science, 17, pp. 98–104. doi: 10.1016/j.cois.2016.08.003.

Stamp, N. E. and Yang, Y. (1996) ‘Response of Insect Herbivores to Multiple Allelochemicals Under Different Thermal Regimes’, Ecology, 77(4), pp. 1088–1102. doi: 10.2307/2265578.

Stearns, S. C. and Kawecki, T. J. (1994) ‘Fitness Sensitivity and the Canalization of Life-History Traits’, Evolution, 48(5), p. 1438. doi: 10.2307/2410238.

Thomas, C. D. et al. (2004) ‘Extinction risk from climate change’, Nature, 427(6970), pp. 145–148. doi: 10.1038/nature02121.

Tournas, V. H. and Katsoudas, E. (2005) ‘Mould and yeast flora in fresh berries, grapes and citrus fruits’, International Journal of Food Microbiology, 105(1), pp. 11–17. doi: 10.1016/j.ijfoodmicro.2005.05.002.

Vianna, B. da S., et al. (2020) ‘Effects of temperature increase on the physiology and behavior of fiddler crabs’, Physiology and Behavior. Elsevier, 215(December 2018), p. 112765. doi: 10.1016/j.physbeh.2019.112765.

Volkoff, H. and Rønnestad, I. (2020) ‘Effects of temperature on feeding and digestive processes in fish’, Temperature, 7(4), pp. 307–320. doi: 10.1080/23328940.2020.1765950.

Wagner, G. P., Booth, G. and Bagheri-Chaichian, H. (1997) ‘A POPULATION GENETIC THEORY OF CANALIZATION’, Evolution, 51(2), pp. 329–347. doi: 10.1111/j.1558-5646.1997.tb02420.x.

Zanco, B. et al. (2021) ‘A dietary sterol trade-off determines lifespan responses to dietary restriction in Drosophila melanogaster females’, eLife, 10, pp. 1–20. doi: 10.7554/eLife.62335.

