## Supplementary Table 1 for "Fluctuating temperatures exacerbate the effects of nutritional stress during development in *Drosophila melanogaster*"

**Supplementary Table 1.** The effects of fluctuating versus constant temperatures on egg-to-adult viability in populations of *D. melanogaster* from Melbourne and Townsville. Data were analysed using a linear model with mixed effects.

|  | Estimate | Standard error | Z value | P value |
| --- | --- | --- | --- | --- |
| Temperature | -0.196 | 0.078 | -2.495 | <0.001 *** |
| Population | -0.080 | 0.107 | -0.749 | 0.005 ** |
| Temperature: population | -0.006 | 0.112 | -0.052 | 0.958 |
