## Supplementary Table 2 for "Fluctuating temperatures exacerbate the effects of nutritional stress during development in *Drosophila melanogaster*"

**Supplementary Table 2.** The effects of fluctuating versus constant temperatures on wing size in populations of *D. melanogaster* from Melbourne and Townsville. Data were analysed using a linear model with mixed effects.

| Variable | Estimate | Standard error | t value | P value |
| --- | --- | --- | --- | --- |
| Temperature | -0.046 | 0.019 | -2.439 | 0.043* |
| Population | -0.077 | 0.018 | -4.191 | <0.001 *** |
| Temperature: population | -0.038 | 0.027 | -1.434 | 0.151 |
