## Supplementary Table 3 for "Fluctuating temperatures exacerbate the effects of nutritional stress during development in *Drosophila melanogaster*"

**Supplementary Table 3.** Effects of carbohydrate (C), protein (P) and their squares, population (Pop), temperature (T), and interaction terms on egg-to-adult viability under three fluctuating temperature regimes (18±5°C, 25±5°C and 28±5°C).

| Variable | LiChisq | Degrees of freedom | P value |
| --- | --- | --- | --- |
| C | 11.7430 | 1 | *** |
| P | 270.388 | 1 | *** |
| C^2^ | 472.326 | 1 | *** |
| P^2^ | 50.978 | 1 | *** |
| Pop | 272.853 | 1 | *** |
| T | 29.537 | 2 | *** |
| C: P | 1203.649 | 1 | *** |
| Pop: T | 94.029 | 2 | *** |
| C: Pop | 19.734 | 1 | *** |
| C: T | 20.709 | 2 | *** |
| P:Pop | 0.1393 | 1 | 0.709 |
| P:T | 2.163 | 2 | 0.340 |
| C:Pop:T | 2.525 | 2 | 0.283 |
| P:Pop:T | 2.979 | 2 | 0.226 |
| C:P:T | 127.531 | 2 | *** |
| C:P:T | 14.780 | 1 | *** |
| C:P:Pop:T | 54.243 | 2 | *** |
