## Supplementary Table 4 for "Fluctuating temperatures exacerbate the effects of nutritional stress during development in *Drosophila melanogaster*"

**Supplementary Table 4.** Output for the angle $\theta_{i,j},$ which is a proxy for nutritional trade-offs with respect to the ratio of nutrients. The estimates of $\theta_{i,j}$ are shown below for each trait response (viability and wing size) between populations (Melbourne versus Townsville) and amongst temperatures (18±5°C, 25±5°C and 28±5°C) (see Morimoto *et al.*, 2023 and methods above).

| Trait |  | Mean | Standard deviation | Lower confidence interval | Upper confidence interval | Population | Temperature |
| --- | --- | --- | --- | --- | --- | --- | --- |
| Viability |  | -3.34 | 8.30 | -19.61 | 12.93 | Melbourne - Melbourne | 25 - 18 |
| Viability |  | 17.39 | 6.07 | 5.49 | 29.30 | Melbourne - Melbourne | 28 – 18 |
| Viability |  | 20.67 | 8.00 | 4.99 | 36.36 | Melbourne – Melbourne | 28 – 25 |
| Viability |  | 3.60 | 8.22 | -12.50 | 19.71 | Townsville – Melbourne | 18 - 25 |
| Viability |  | -17.13 | 5.81 | -28.63 | -5.62 | Townsville – Melbourne | 18 – 28 |
| Viability |  | -10.43 | 6.06 | -22.31 | 1.45 | Townsville – Melbourne | 25 - 28 |
| Viability |  | 6.69 | 6.29 | -5.65 | 19.02 | Townsville - Townsville | 25 - 18 |
| Viability |  | 3.37 | 5.80 | -8.00 | 14.73 | Townsville – Townsville | 28 – 18 |
| Viability |  | -3.34 | 6.01 | -15.12 | 8.44 | Townsville – Townsville | 28 – 25 |
| Wing size |  | -17.15 | 8.40 | -33.62 | -0.69 | Melbourne - Melbourne | 25 - 28 |
| Wing size |  | -30.64 | 6.49 | -43.36 | -17.93 | Melbourne-Melbourne | 18 - 25 |
| Wing size |  | -13.58 | 6.40 | -26.12 | -1.03 | Melbourne – Melbourne | 18 - 25 |
| Wing size |  | 0.01 | 10.82 | -21.20 | 21.20 | Townsville - Melbourne | 25 - 28 |
| Wing size |  | -30.32 | 6.57 | -43.20 | -17.44 | Townsville – Melbourne | 18-28 |
| Wing size |  | -13.25 | 6.45 | -25.90 | -0.60 | Townsville - Melbourne | 18-25 |
| Wing size |  | 0.68 | 10.936 | -20.76 | 22.12 | Townsville - Townsville | 25-28 |
| Wing size |  | -29.65 | 6.82 | -43.55 | -12.17 | Townsville – Townsville | 18-28 |
| Wing size |  | -30.36 | 9.28 | -48.55 | -12.17 | Townsville - Townsville | 18 – 25 |
