## Supplementary Table 5 for "Fluctuating temperatures exacerbate the effects of nutritional stress during development in *Drosophila melanogaster*"

**Supplementary Table 5.** Effects of carbohydrate (C), protein (P), population (Pop), temperature (T), and interaction terms on female wing centroid size under three fluctuating temperature regimes (18±5°C, 25±5°C and 28±5°C).

| Variable | LiChisq | Degrees of freedom | P value |
| --- | --- | --- | --- |
| C | 0.847 | 1 | 0.358 |
| P | 68.559 | 1 | *** |
| C^2^ | 62.671 | 1 | *** |
| P^2^ | 39.654 | 1 | *** |
| Pop | 1022.825 | 1 | *** |
| T | 7093.825 | 2 | *** |
| C: P | 0.0625 | 1 | 0.803 |
| Pop: T | 6.673 | 2 | 0.083 |
| C: Pop | 1.823 | 1 | 0.178 |
| C: T | 38.106 | 2 | *** |
| P:Pop | 7.332 | 1 | ** |
| P:T | 180.112 | 2 | *** |
| C:Pop:T | 6.236 | 2 | * |
| P:Pop:T | 2.377 | 2 | 0.305 |
| C:P:T | 64.541 | 2 | *** |
| C:P:T | 8.438 | 1 | ** |
| C:P:Pop:T | 6.905 | 2 | * |
